# Maternal IBD-related antibodies are associated with early life gut inflammatory status and microbiota composition: insights from cord blood

**DOI:** 10.1101/2025.08.04.668517

**Authors:** Taegyu Kim, Paola Nicoletti, Maria Manuela Estevinho, Leonid Tarassishin, Mellissa Picker, Anketse Debebe, Isabella Nguyen, Rhiannon Dutra-Clarke, Courtney Young, Joanne Stone, Ariella Bar-Gil Shitrit, Manasi Agrawal, Jean-Frederic Colombel, Joana Torres, Siew C Ng, Lin Zhang, Dermot P. B. McGovern, Inga Peter

**Affiliations:** Department of Genetics and Genomic Sciences, Icahn School of Medicine at Mount Sinai, New York, NY, USA; Department of Gastroenterology, Unidade Local de Saúde Gaia Espinho, Vila Nova de Gaia, Portugal; Department of Biomedicine, Unit of Pharmacology and Therapeutics, Faculty of Medicine, University of Porto, Porto, Portugal; F. Widjaja Inflammatory Bowel Disease Institute, Cedars-Sinai Medical Center, Los Angeles, CA, USA; Division of Maternal Fetal Medicine, Department of Obstetrics, Gynecology and Reproductive Science, Icahn School of Medicine at Mount Sinai, New York, NY, USA; Digestive Diseases Institute, The Hebrew University of Jerusalem, Shaare Zedek Medical Center, Jerusalem, Israel; The Henry D. Janowitz Division of Gastroenterology, Department of Medicine, Icahn School of Medicine at Mount Sinai, New York, NY, USA; Division of Gastroenterology, Hospital da Luz & Hospital Beatriz Ângelo, Portugal; Microbiota I-Center (MagIC), Hong Kong SAR, China; Department of Medicine and Therapeutics, New Cornerstone Science Laboratory, The Chinese University of Hong Kong, Hong Kong SAR, China; State Key Laboratory of Digestive Diseases, Li Ka Shing Institute of Health Sciences, The Chinese University of Hong Kong, Hong Kong SAR, China; Department of Anaesthesia and Intensive Care, The Chinese University of Hong Kong, Hong Kong SAR, China

**Keywords:** anti-CBir1, ANCA, anti-OmpC, ASCA, inflammatory bowel disease, pregnancy

## Abstract

**Purpose:** Antibodies in peripheral blood are used to aid in the diagnosis of inflammatory bowel disease (IBD), but their presence in neonatal cord blood and potential effects on early life development remain unknown.

**Methods:** We measured anti-CBir1, ANCA, anti-OmpC, ASCA IgA, and ASCA IgG levels in the cord blood of babies born to 78 mothers with or without IBD. Their association with fecal calprotectin (FC), and microbiota composition, characterized by 16S rRNA sequencing, was assessed throughout pregnancy and during the first 3 years of life using linear mixed-effects models.

**Results:** Antibodies were detected in cord blood, with significantly higher levels of anti-CBir1 and ASCA IgG in babies born to mothers with Crohn’s disease (p = 0.002) and higher abundance of ANCA and anti-OmpC in babies of mothers with ulcerative colitis (p = 0.002), compared to controls. ASCA IgG levels positively correlated with babies’ FC (p = 0.006), while babies’ microbiota Shannon diversity was negatively associated with ANCA, anti-OmpC, and ASCA IgA levels (p = 0.003, 0.04, and 0.008, respectively). *Romboutsia* spp., *Citrobacter* spp., *Pseudomonas* spp., *Clostridiaceae*, *Clostridia*, and *Varibaculum* spp. were positively correlated with either or both ANCA and anti-OmpC levels (all multiple testing adjusted q < 0.1).

**Conclusion:** Our findings suggest that maternal IBD-associated antibodies cross the placenta barrier and may be associated with intestinal inflammation and imbalanced microbiota colonization. Whether these serological profiles negatively influence the priming of the baby’s immune system or IBD risk later in life remains to be determined.

## Introduction

The maternal immune health and gut microbiota composition have been shown to play a crucial role in the offspring’s immune system development both in utero and after birth ^1^. During pregnancy, the immune-modulating factors – including maternal antibodies and cytokines – are transmitted from the mother to the fetus via the placenta and cord blood. Maternal antibodies provide passive immunity and immediate protection, while cytokines influence how neonatal immune cells are primed to respond to future immune challenges ^1–3^, such as infections and allergens ^4,5^. At birth, vaginal seeding is recognized as an essential mechanism through which newborns acquire initial microbial colonization ^6^. However, by four months of age, most gut colonizers are sourced from the maternal gut microbiome rather than vaginal flora ^7^. The mother-baby immune-related interactions continue with breastfeeding, which facilitates the transfer of gut microbiome-induced maternal antibodies to the neonatal intestine, along with secretory IgA and other bioactive factors ^8–10^.

Therefore, maternal pre-existing health conditions can affect the infant’s immune system development, mediating long-lasting health outcomes ^11^, including the predisposition to immune-mediated diseases later in life ^12,13^. Our team has shown that babies born to mothers with inflammatory bowel disease (IBD), a chronic inflammatory condition of the gastrointestinal tract that may be caused by loss of mucosal tolerance to the commensal microbiota in genetically susceptible individuals ^14,15^, have lower gut bacterial diversity, resulting in abnormal imprinting of the intestinal immune system in germ-free mice colonized with these infants’ stool ^16^. Moreover, we demonstrated higher levels of fecal calprotectin (FC) - a well-established biomarker of the presence and degree of intestinal inflammation ^17^ and a predictor of future IBD risk ^18^ - in unaffected offspring of IBD mothers up to at least 3 years of age ^19^. These findings suggest that maternal IBD may negatively affect fetal-maternal interactions.

Serological antibodies in the peripheral blood, such as anti-flagellin CBir1 (anti-CBir1), anti-outer membrane porin C (anti-OmpC), anti-neutrophil cytoplasmic antibody (ANCA), and anti-*Saccharomyces cerevisiae* antibody (ASCA), recognize a range of antigens, including microbial proteins (e.g., CBir1, OmpC), host-derived cytoplasmic proteins (e.g., ANCA), and oligosaccharide antigens (e.g., ASCA).

These antibodies have been used to aid in the differential diagnosis of IBD and its subtypes, Crohn’s disease (CD) and ulcerative colitis (UC) ^20–22^. High levels of anti-CBir1 indicate a distinct immune response in CD ^23^. ANCA antibodies are more prevalent in UC, being detected in 60–80% of UC patients but only in 5–20% of CD patients ^24–26^. ASCA is more frequently detected in CD than UC ^27,28^. A combination of positive ASCA IgA and IgG with negative ANCA is strongly associated with CD and has prognostic value for disease progression ^29,30^. While these markers aid in differentiating IBD from non-IBD and CD from UC, their transmission from mother to fetus and their impact on the factors driving immune development during early life remain poorly explored.

Cord blood provides a unique opportunity to study the direct maternal transfer of IBD-associated antibodies to the baby, which can potentially influence immune development, modulate gut microbiome colonization, and impact inflammatory pathways during early life, a critical period for establishing long-term immune homeostasis ^31^. A recent study established that antibodies like ASCA are not just passive markers but are active initiators of the inflammatory cascade ^32^, raising a question of what microbial triggers might be responsible for producing these pathogenic antibodies. Therefore, the study aimed to investigate the levels of the top IBD-associated antibodies (anti-CBir1, anti-OmpC, ANCA, and ASCA) in cord blood from pregnancies of women with and without IBD and determine their association with the intestinal inflammatory status and gut microbiota composition of both mother and infant to identify potential relationships that may influence early gut colonization. A better understanding of feto-maternal interactions may help shed light on the potential pathways that promote the risk transmission of immune-mediated diseases, such as IBD, and provide valuable insights to inform early intervention strategies and promote long-term health.

## Material and methods

### Participant characteristics

This study was conducted with 17 UC, 11 CD, and 50 control pregnant women in a subset of the MECONIUM (Exploring *Mec*hanisms of Disease Transmissi*on I*n *U*tero through the *M*icrobiome) cohort recruited between 2015 and 2021. Among them, 4 UC and 22 control women were recruited from Hong Kong (HK) via the medical and IBD clinics at the Prince of Wales Hospital, while others were recruited in the US.

The information about age at enrollment, race/ethnicity, smoking status, baseline body mass index (BMI), and IBD-related drug usage during pregnancy (e.g., aminosalicylates, steroids, immunomodulators [IMM], and anti-tumor necrosis factor [anti-TNF-α]) was collected for mothers.

Disease activity was assessed prospectively by Physician Global Assessment and evaluated as either active or in remission during each trimester. Harvey–Bradshaw Index (HBI) and the Mayo score (without an endoscopic score) were used to assess disease activity in CD and UC, respectively. CD was considered in remission if HBI was < 5, while UC was in remission if the partial Mayo score was < 2.

In the offspring, the collected data included gestational age at delivery, delivery mode (vaginal or cesarean), sex, exposure to antibiotics, and feeding behavior (exclusive breastfeeding, exclusive formula feeding, or mixed feeding). Infant feeding practices were surveyed at one week, two weeks, one month, two months, and three months postpartum using two dichotomous (yes/no) questions: ‘Was your child breastfed during this period?’ and ‘Did you breastfeed exclusively during this period?’

This study was approved by the Institutional Review Board of the Icahn School of Medicine at Mount Sinai (HS#14-00554) and Joint Chinese University of Hong Kong - New Territories East Cluster Clinical Research Ethics Committee (CREC no. 2016.547).

### Cord blood sample collection and antibody response analysis

Venous umbilical cord blood was collected at the time of delivery using a non-EDTA blood collection tube for the US samples. Umbilical cord blood (for buffy coat, serum, and plasma [EDTA]) was also collected at the HK site. The tubes were left standing for 30 minutes to allow clotting, and then centrifuged at 1520 × *g* for 15 minutes at 4°C to collect the serum samples. Serum samples were stored at -20°C until the analysis. HK samples were transported in liquid nitrogen to the US for further processing and analysis.

Patient sera were shipped on ice using a next-day air service and analyzed in a blinded manner at Cedars-Sinai Medical Center for the expression of several antibodies by enzyme-linked immunosorbent assay (ELISA) following previously described protocols ^33–37^. The analysis measured both ASCA IgG and ASCA IgA, as well as ANCA, anti-OmpC, and anti-CBir1 (for which IgG and IgA isotypes were not separately measured). In short, test sera were analyzed alongside control reference sera from Cedars-Sinai’s MIRIAD Biobank, including negative and positive controls. ANCA expression was assessed by fixed neutrophil ELISA with methanol-fixed human neutrophils, while ASCA, anti-OmpC, and anti-CBir1 expressions were determined by reactivity to *Saccharomyces cerevisiae mannan*, *E. coli* OmpC, and CBir1 flagellin, respectively, on high-binding polystyrene microtiter plates. The assays involved antigen coating, blocking with 0.5% BSA/PBS, incubation with diluted sera, and detection with specific secondary antibodies, followed by pNPP substrate. Plates were read on an Emax microplate reader (Molecular Devices, CA, USA) when positive control optical density (OD) reached 1.00–2.00 OD above blanks. Raw OD values were normalized to ELISA units (EU) relative to positive controls (100 EU), with seropositivity defined by assay-specific cutoffs (ANCA >30 EU, ASCA IgA >20 EU, ASCA IgG >40 EU, anti-OmpC >23 EU, anti-CBir1 >25 EU), which were established with peripheral blood during assay development. These thresholds were typically defined as the mean plus two standard deviations of seroreactivity scores from a cohort of clinically well-characterized CD and UC patient sera for each assay.

### Fecal calprotectin (FC) analysis

During pregnancy, stool samples were collected at home with an ALPCO Diagnostics EasySampler Stool Collection kit (ALPCO Life Science, Salem, NH, USA) at the 1^st^ (T1), 2^nd^ (T2), and 3^rd^ (T3) trimesters. Following delivery, the babies’ diapers or stool samples were collected at 1W (week), 2W, 1M (month), 2M, 3M, 1Y (year), 1.5Y, 2Y, 3Y, 4Y, and 5Y. Samples were shipped on ice to the lab and stored at -20°C or -80°C until further analysis. The FC concentration from both mothers and babies was determined using a CALPROLAB Calprotectin ELISA kit (Svar Life Science, Lysaker, Norway), as described elsewhere ^19^. For FC, data up to 3 years of age were used.

### Stool microbiome analysis

A subset of stool samples (maternal samples n = 108, baby samples n = 401) were profiled using bacterial 16S rRNA gene sequencing with HiSeq paired-end 250×2 fast mode (Illumina, San Diego, CA, USA). The fecal DNA extraction and dual-barcoded 16S V4 PCR amplicon preparation was performed as previously described ^16^. For sequence analysis, *mothur* was employed ^38^ using a protocol similar to the MiSeq SOP ^39^. In short, paired-end reads were merged, and sequences with one or more ambiguous bases or a homopolymer longer than eight nucleotides were removed. Sequences were aligned to the SILVA v.138 reference file ^40^, and chimeric sequences were then identified and discarded using VSEARCH ^41^. A Bayesian classifier and the SILVA v.138 reference database were used to classify the sequences ^42^. Sequences that did not classify as *Bacteria* or *Archaea* were discarded. The number of sequences was normalized before sequence clustering into operational taxonomic units (OTUs) at a 3% similarity threshold. Then, the numbers of OTUs and Shannon diversity index were calculated for the microbiota alpha diversity, and Bray-Curtis dissimilarity was calculated for the microbiota beta diversity.

### Statistical analysis

Fisher’s exact test and Wilcoxon rank-sum test were employed to evaluate statistical differences between any two groups. Principal Component Analysis (PCA) was conducted to visualize the variation in serological marker levels across the samples, along with the loadings of each serological marker on the principal components. Multiple logistic regression analyses were performed to generate receiver operating characteristic (ROC) curves to assess predictive variables for maternal IBD diagnoses. All statistical analyses were performed using GraphPad Prism, version 10.2.3 (GraphPad, San Diego, CA, USA).

To assess the significant differences in serological marker levels between groups of samples, an analysis of similarities (ANOSIM) was performed using the R package *vegan* ^43^, and confidence ellipses were calculated by the R package *MASS* ^44^. To account for repeated measures, a linear mixed-effects model was employed using the R package *lme4* ^45^, with subject ID as a random effect and fixed effects drawn from disease type, time point, BMI, anti-TNF-α usage, and geographic location. FC values were log-transformed prior to analyses that assume linear relationships, including Pearson’s correlation and linear mixed-effects model analysis. Principal coordinates analysis (PCoA) and analysis of molecular variance (AMOVA) were conducted using *mothur* ^38^. Multiple testing correction using false discovery rate was applied with a significance threshold of q < 0.1 ^46^.

## Results

### Participant characteristics

The characteristics of the 17 UC, 11 CD, and 50 control subjects are summarized in **Table 1**.

**Table 1.** Participant characteristics

|  | UC, N = 17 | CD, N = 11 | Control, N = 50 | p, UC vs. CD<sup>3</sup> | p, UC vs. control | p, CD vs. control |
| --- | --- | --- | --- | --- | --- | --- |
| **Maternal characteristics** |  |  |  |  |  |  |
| Age at enrollment (years) | 34 (33, 37)<sup>1</sup> | 34 (31, 35) | 34 (32, 37) | 0.29 | 0.56 | 0.44 |
| Jewish ancestry (Yes) | 5/15 (33%)<sup>2</sup> | 6/11 (55%) | 6/49 (12%) | 0.43 | 0.11 | **<u>0.01</u>** |
| Ex-smoker (Yes) | 2/16 (13%) | 0/11 (0%) | 5/49 (10%) | 0.50 | > 0.99 | 0.57 |
| Body mass index | 24.4 (22.2, 25.7) | 24.6 (22.5, 29.0) | 25.3 (23.1, 27.1) | 0.50 | 0.38 | 0.79 |
| *Drug usage during pregnancy* |  |  |  |  |  |  |
| Aminosalicylates | 12/16 (75%) | 5/10 (50%) | - | 0.23 | - | - |
| Steroids | 1/16 (6%) | 1/10 (10%) | - | > 0.99 | - | - |
| IMM | 3/16 (19%) | 1/10 (10%) | - | > 0.99 | - | - |
| Anti-TNF-α | 5/16 (31%) | 7/10 (70%) | - | 0.11 | - | - |
| **Infant characteristics** |  |  |  |  |  |  |
| Gestational age at delivery, weeks | 39.0 (38.8, 39.2) | 38.8 (37.4, 39.4) | 39.1 (38.4, 40.0) | 0.71 | 0.64 | 0.26 |
| Sex (male : female) | 9:7 (56:44%) | 7:4 (64:36%) | 24:23 (51:49%) | > 0.99 | 0.78 | 0.52 |
| Delivery mode (vaginal : cesarean) | 8:8 (50:50%) | 3:8 (27:73%) | 32:15 (68:32%) | 0.43 | 0.24 | **<u>0.02</u>** |
| Antibiotic prior to delivery | 10/14 (71%) | 9/11 (82%) | 19/32 (59%) | 0.66 | 0.52 | 0.28 |
| Exposure to antibiotics in hospital | 1/12 (8%) | 2/10 (20%) | 3/28 (11%) | 0.57 | > 0.99 | 0.59 |
| Exclusive breastfeeding in the first 90 days | 4/15 (27%) | 1/11 (9%) | 10/41 (24%) | 0.36 | > 0.99 | 0.42 |

<sup>1</sup>median with interquartile range interval
<sup>2</sup>based on non-missing data
<sup>3</sup>p-values from group comparisons (Fisher's exact test for categorical values and the Wilcoxon rank-sum test for continuous values); p < 0.05 are shown in bold and underlined

Aside from a higher proportion of mothers in the CD group reporting Jewish ancestry compared to controls (55% vs. 12%, p = 0.01), no other significant differences were observed between the groups (p > 0.05).

Infant sex, exposure to antibiotics in the hospital, and feeding behavior up to 90 days did not differ significantly between groups, although a higher proportion of CD mothers underwent cesarean delivery compared to controls (73% vs. 32%, p = 0.02).

### Serological marker levels in cord blood

Serological antibodies were detectable in cord blood samples (**Figure 1**). With the standard seropositivity threshold established for peripheral blood during assay development at the laboratory, anti-CBir1 was positive in 0% of UC patients, 27% of CD patients, and 2% of controls, while ANCA was positive in 29% of UC patients, 9% of CD patients, and 2% of controls (**Table 2**). The seropositivity rates of the UC group were distinct from the CD and the control groups (both p < 0.0001); however, the CD group was not distinct from the control group (p = 0.29). This suggests that the standard peripheral blood positivity cutoff may be overly stringent for cord blood samples, and the placental transmissivity of ASCA is lower than that of anti-CBir1 and ANCA. Indeed, when the analysis was repeated using a 50% lower cutoff threshold, a provisional threshold selected due to the absence of established reference values in cord blood, a significant difference emerged between the CD and control groups (p = 0.0006), and all inter-group comparisons became statistically significant.

**Figure 1.**
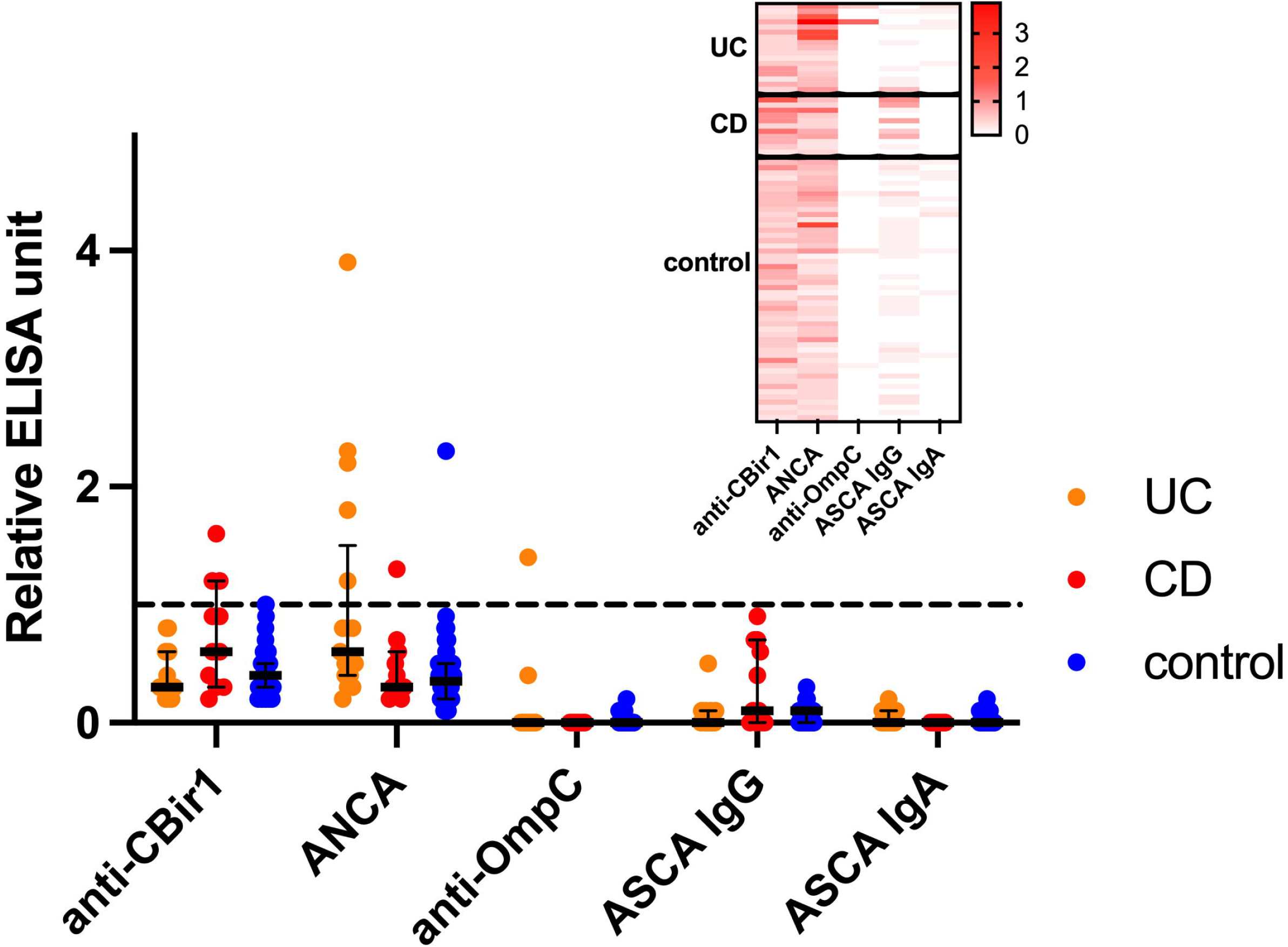
Relative ELISA units of five serological biomarker levels in cord blood with respect to the seropositivity threshold used in peripheral blood. The threshold used in peripheral blood is shown as a black dashed line. A heatmap shows the relative ELISA units of each biomarker in each individual.

**Table 2.**
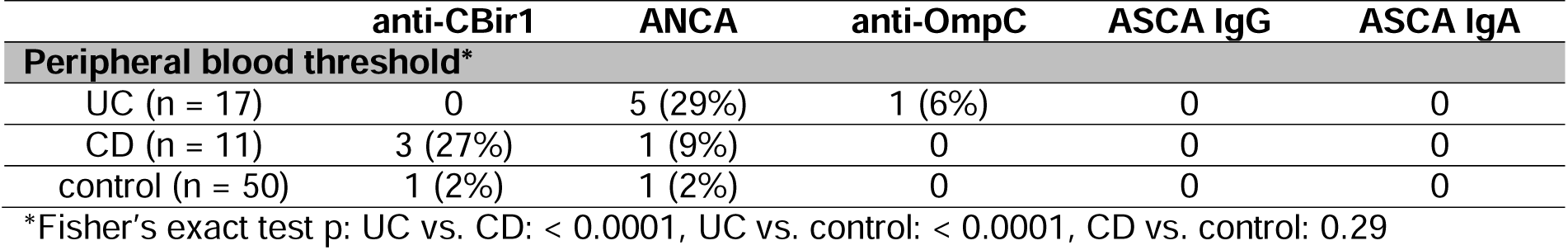
Number of seropositivity cases and percentages in parentheses in each group

Anti-OmpC and ASCA IgA were not detected or detected at a very low level in most samples. Anti-OmpC was positive based on the standard seropositivity threshold in one UC sample. ASCA IgA was detected in 10 out of 17 UC samples, 4 out of 11 CD samples, and 15 out of 50 control samples at a level lower than the threshold. The detected signals were weaker compared to ASCA IgG, which was present in all cord blood samples.

### Maternal disease classification

Serological marker abundances in cord blood samples were distinctive based on the maternal IBD status (**Figure 2A**; ANOSIM test for separation between UC vs. CD [p = 0.025], UC vs. control [p = 0.002], and CD vs. control [p = 0.002]). ASCA IgG and anti-CBir1 drove the differences with CD, while anti-OmpC and ANCA levels differentiated the UC group from others.

**Figure 2.**
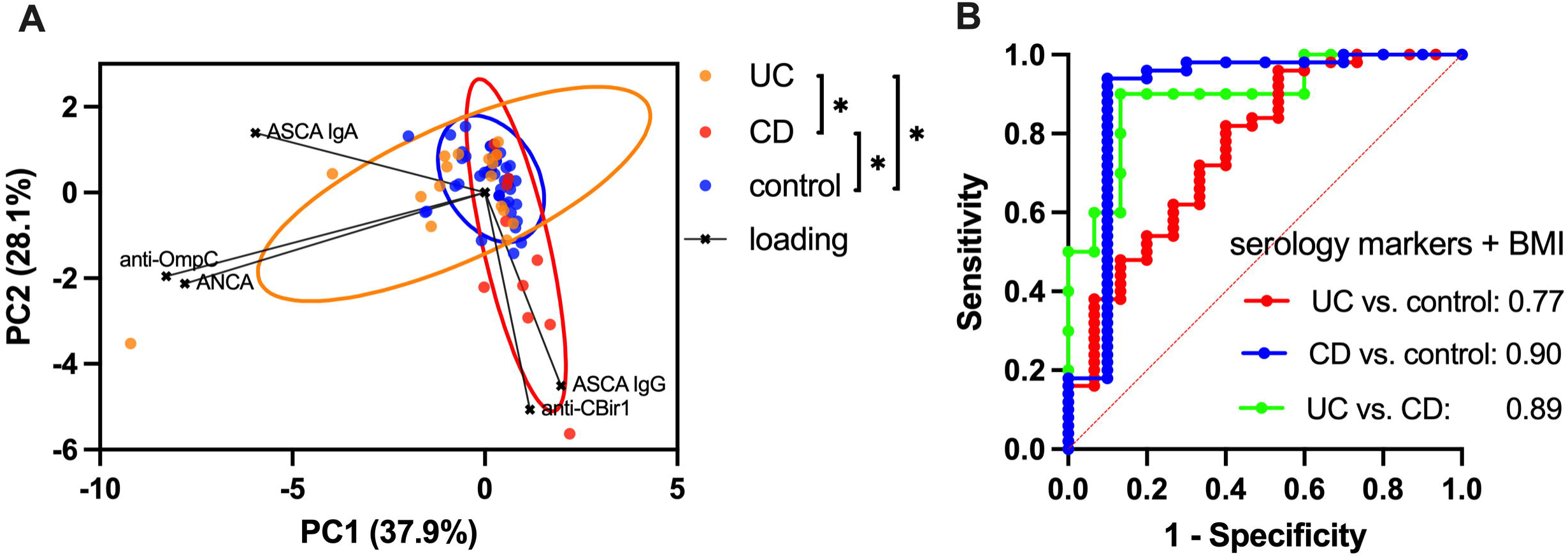
(A) Principal component analysis (PCA) of cord blood serological marker levels; asterisks (*) indicate significant group differences as determined by analysis of similarity (ANOSIM; p < 0.05) and 90% confidence ellipses are shown, (B) Receiver operating characteristic (ROC) curves for different disease types (UC, ulcerative colitis; CD, Crohn’s disease, and control), derived from logistic regression models using serology marker levels and maternal BMI. The areas under the curves (AUCs) are displayed in each plot. Detailed test results with sensitivity, specificity, negative/positive predictive values, and accuracy with and without BMI are shown in **Table S1**.

Logistic regression analysis also showed that those markers can have predictive value for maternal disease subtypes (**Figure 2B**). Sensitivity for detecting true UC and CD cases compared to control groups was 29.4% and 54.6%, respectively, while specificity was high for both at 98.0%, with corresponding area under the curves (AUCs) of 0.77 and 0.90 (**Table S1**). In all three group comparisons among UC, CD, and control, the negative predictive value, positive predictive value, and overall accuracy were all equal to or greater than 80.0%. When the logistic regression model included the maternal BMI, to account for possible hemodilution of serologic markers, sensitivity for detecting true UC and CD cases compared to the control group increased to 33.3% and 70.0%, while specificity remained the same at 98.0% (**Table S1**).

Within IBD mothers, those treated with any IBD-directed drug exhibited higher cord blood marker levels overall, particularly anti-TNF-α (**Figure S1**). In the case of UC, the anti-TNF-α-treated group showed significantly higher ANCA levels (p = 0.02). For CD, the anti-TNF-α-treated group demonstrated elevated anti-CBir1 levels with borderline statistical significance (p = 0.08). The median ANCA level appeared to be higher in UC mothers with active disease during pregnancy, although the difference was not statistically significant (p = 0.50), likely due to the limited sample size (5 active, 8 inactive, 4 unknown; **Figure S1**).

Maternal BMI measured in early pregnancy among IBD mothers was negatively correlated with all serological biomarker levels in cord blood, both in UC and CD groups (rho ranged from -0.16 to -0.68), with the statistical significance shown between BMI and anti-OmpC in the CD group (p = 0.04) (**Figure S2**). The correlation between BMI and serological biomarker levels in the control group was not statistically significant (rho ranged from -0.02 to 0.27, all p > 0.06; **Figure S2**).

### Fecal calprotectin (FC) and serological markers

Cord blood serology marker levels showed weak positive correlations with maternal FC levels during pregnancy, except for ASCA IgA (**Figure 3**). Although individual serological markers were not significantly associated with maternal FC levels, a composite z-score combining all five markers was significantly associated with maternal FC after adjusting for time in a multiple linear regression model (p = 0.019). In the baby samples, there was a significant overall association between ASCA IgG and FC levels, after adjustment for time point (p = 0.006), and after adjustment for time point and maternal IBD status (p = 0.05). Pearson’s correlation analysis between ASCA IgG and FC levels at individual time points further supported a positive relationship, with statistically significant associations detected at 2 weeks, 3 months, and 1 year (p = 0.04, 0.04, and 0.02, respectively; **Figure 3**).

**Figure 3.**
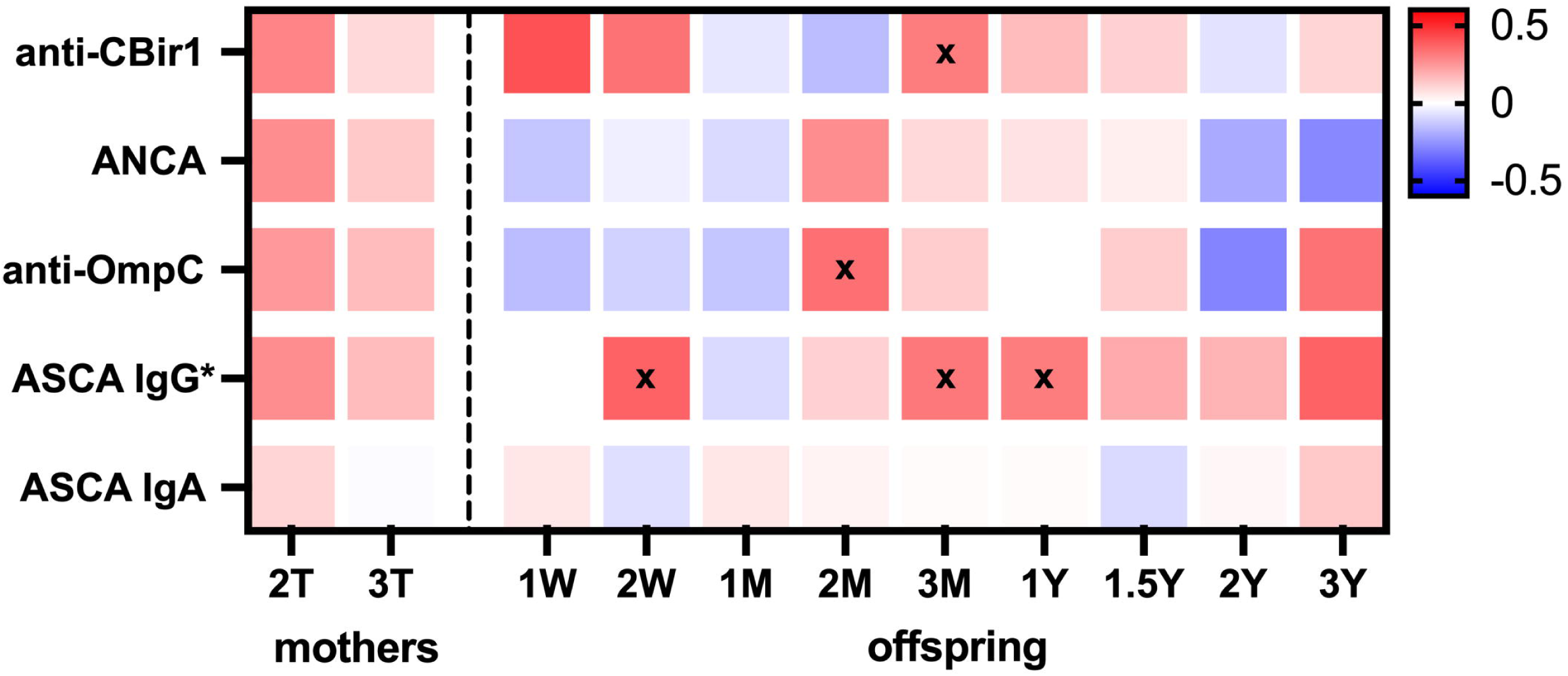
Pearson’s correlation coefficient heatmap between log-transformed fecal calprotectin levels and cord blood serological markers in mothers (2nd trimester [2T] and 3rd trimester [3T]) and their babies at multiple time points (W: weeks, M: months, Y: years). An asterisk (*) indicates a statistically significant association in baby samples across multiple time points, based on a mixed-effects model adjusted for time and maternal disease status. ‘x’ marks denote time points with statistically significant positive correlations (plJ<lJ0.05).

### Stool microbiome and taxa abundance

To assess associations between cord blood serological biomarker levels and maternal stool microbiota alpha diversity or taxa abundance throughout pregnancy, linear mixed-effects models were used while adjusting for disease subtype, time point, BMI, anti-TNF-α therapy, and geographic location (**Figure 4A**). For taxa abundance, the analysis included the top 105 bacterial taxa with an average relative abundance >0.5% and revealed a significant negative association between anti-CBir1 antibody levels and the following taxa: *Streptococcus* spp., *Romboutsia* spp., *Enterobacteriaceae*, *Actinomyces* spp., *Granulicatella* spp., and *Lactobacillales*. Conversely, anti-CBir1 levels showed a positive association with *Butyricimonas* spp., *Bacteroidales*, *Hungatella* spp., *Ruminococcaceae*, *Tannerellaceae*, *Ruminococcaceae* UBA1819, and *Parabacteroides* spp. (all q < 0.1). Additionally, ANCA levels were positively associated with the number of observed OTUs (p = 0.03) and with *Barnesiella* spp. abundance (q = 0.01).

**Figure 4.**
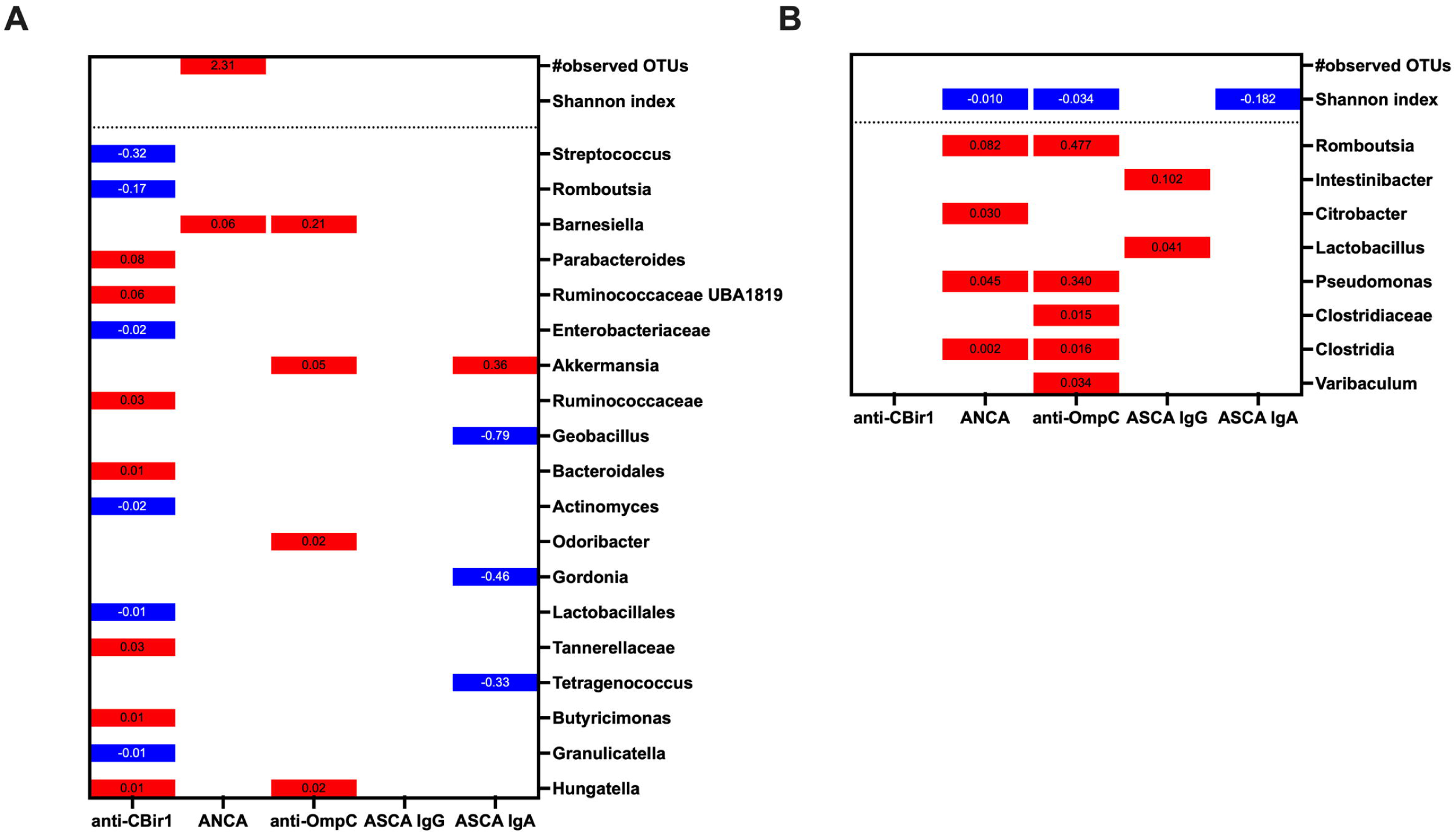
Sparse plots showing statistically significant beta coefficients from linear mixed-effects models assessing associations between cord blood serological biomarker levels and (**A**) maternal stool alpha diversity or taxa abundance, adjusted for disease type, time point, BMI, anti-TNF-α usage, and geographic location; and (**B**) baby stool alpha diversity or taxa abundance, adjusted for time point and geographic location. The red columns show positive associations, and blue columns show negative associations. For alpha diversity, the results with p < 0.05 are shown. Taxa with an average abundance >0.5% were tested (105 for mothers and 92 for babies), and results are shown only for associations with a multiple testing-adjusted q < 0.1.

In babies, we examined the association between cord blood serological biomarkers and gut microbiome alpha diversity and the relative abundance of the top 92 bacterial taxa with an average relative abundance >0.5%, while adjusting for time point and geographic location (**Figure 4B**). Shannon diversity index was negatively associated with levels of ANCA, anti-OmpC, and ASCA IgA (p = 0.003, 0.04, and 0.008, respectively). Among individual bacterial taxa, *Intestinibacter* spp. and *Lactobacillus* spp. were positively associated with ASCA IgG levels (q = 0.07 and 0.03, respectively). *Romboutsia* spp., *Citrobacter* spp., *Pseudomonas* spp., *Clostridiaceae*, *Clostridia*, and *Varibaculum* spp. were positively associated with either or both ANCA and anti-OmpC levels (all q < 0.1).

To compare cord blood markers with gut microbiota alpha and beta diversity, we focused on anti-CBir1 and ANCA, the two most frequently detected antibodies. Subjects were stratified into four groups based on high versus low levels of each marker to enable group-level comparisons. The 50% of standard peripheral blood seropositivity cutoff was used to define high, resulting in 25 subjects in the high-anti-CBir1 group (including 7 of the 11 CD subjects) and 25 subjects in the high-ANCA group (including 12 of the 17 UC subjects). Maternal stool microbiota composition (beta diversity) significantly differed between the high-ANCA group and the both-low group, both at 2T and 3T, based on the AMOVA test (p = 0.012 and 0.031, respectively; **Figure S3**). Also, the high-ANCA group showed a higher number of observed OTUs (alpha diversity) compared to the both-low group at T3, although the difference was borderline significant (p = 0.053; **Figure S4**).

The baby stool microbiota showed significant differences in both alpha and beta diversity between the high-ANCA group and the both-low group at 1, 2, and 3 months of age. AMOVA testing revealed significantly different microbial community composition between the two groups at these time points (p = 0.004, <0.001, and 0.003, respectively; **Figure S5**). However, by 1 year of age, the difference was no longer significant (F = 1.2, p = 0.23). Similarly, Shannon diversity was significantly lower in the high-ANCA group compared to the both-low group at 1, 2, and 3 months (p = 0.003, 0.006, and 0.016, respectively; **Figure S6A**), but this difference was not observed after 1 year (**Figure S6B**).

## Discussion

Our study demonstrates, for the first time, that IBD-associated microbial antibodies - predominantly IgG isotypes such as ASCA, anti-CBir1, ANCA, and anti-OmpC - are detectable in cord blood, with higher levels observed in offspring of women living with IBD. Antibody profiles also differ according to maternal disease type, distinguishing between UC and CD. Serological antibody levels showed a positive relationship with maternal FC levels during pregnancy, whereas ASCA IgG levels were positively correlated with the offspring FC levels throughout first months of life. In addition, anti-CBir1, ANCA, and anti-OmpC levels were associated with taxa abundances in the gut microbiota of mothers and their offspring. Our results suggest that pregnant women with IBD have elevated biomarkers of immune response due to mucosal barrier dysfunction and microbial translocation, passing them to their babies, which may influence intestinal health and bacterial diversity and shape early life immune system development.

Cord blood is a readily accessible sample obtained at birth that can be a valuable resource for risk stratification. Previous studies have identified specific cytokine-related markers that were predictive of subsequent bronchopulmonary dysplasia development in premature infants ^47^ or durability of infant vaccine responses ^48^. Maternal IgG antibodies cross the placenta starting in the second trimester of gestation and can be detected in infants up to 6 months of age, while IgA antibodies cross the placenta only negligibly due to their lack of binding to the neonatal Fc receptor and their larger size, particularly in the dimeric form ^5,49,50^. In our study, antibody profiles were predictive of maternal IBD diagnosis and differed by IBD type: ASCA IgG and anti-CBir1 were elevated in the CD group, while anti-OmpC and ANCA were higher in the UC group. This is in line with the reports in the peripheral blood showing elevated ASCA and anti-CBir1 in CD patients, and ANCA in UC patients ^23–28^. However, the positivity of anti-OmpC has been shown to be more frequent in CD (30-60%) compared to UC (5-24%) ^51,52^, whereas in the cord blood, we detected them exclusively in mothers with UC, a finding that warrants further validation and raises the question of potential disease-specific placental transfer mechanisms.

Alongside IBD diagnosis, maternal medication, particularly anti-TNF-α therapy, and BMI during pregnancy seemed to be related to cord blood antibody transmission. Notably, we showed that treatment with TNF-α inhibitors was associated with increased levels of several serological biomarkers, with a significant difference observed in the ANCA levels between treated and untreated UC patients. While TNF-α inhibitors are typically used to suppress inflammation, our finding of elevated cord blood ANCA levels in the anti-TNF-α-treated group may reflect paradoxical immune activation, or an induced autoantibody response, as has been observed in other TNF-induced autoimmune phenomena ^53^. Alternatively, these findings may reflect underlying disease severity not fully controlled by TNF-α inhibitors.

While TNF-α inhibitors have been shown to be transferred across the placenta and be detectable in infants up to 6 months after birth ^49^, it remains to be determined whether these drugs directly suppress the infant’s own immune system ^54^ or indirectly alter it by increasing the amount of maternal antibodies transferred across the placenta. In contrast, maternal baseline BMI showed a trend towards a negative relationship with biomarker levels, which could be attributed to physiological changes in pregnancy, including hemodilution. While the direct relationship between BMI and IBD-associated antibody levels during pregnancy has not been reported, one study observed a higher, though not statistically significant, BMI and obesity rate in the low anti-CBir1 group compared to the high anti-CBir1 group ^55^. These observations emphasize the importance of considering the role of both disease-related and maternal physiological factors in immune “inheritance.”

Maternal gut inflammation may also affect the antibody transmission to the baby by modulating placental barrier permeability or Fc receptor expression, thereby enhancing antibody transfer. FC is a well-established intestinal inflammatory marker and a marker of subclinical immune activation ^19^. Our study showed that maternal FC levels during pregnancy were positively associated with cord blood antibody levels, with a similar relationship reported in the peripheral blood ^56^. Subsequently, cord blood ASCA IgG levels were also positively correlated with infant FC levels, suggesting that maternal gut inflammation may have an effect on neonatal gut health, which in turn may contribute to the priming of the immune system and susceptibility to immune-mediated diseases later in life ^57^. Further studies are needed to elucidate the mechanistic pathways between maternal ASCA IgG and future disease susceptibility in the baby.

Microbial taxa can have a bidirectional association with antibody levels. Microbes shape immunity by stimulating antibody response; conversely, the antibodies can influence microbes by targeting bacteria for immune clearance. Cord blood may facilitate the communication between the maternal and neonatal gut microbiota, reflecting shared immunological and microbial exposure, and consequently playing a role in shaping the development of the early life microbiome. Previous studies have shown that prenatal inflammatory signals can influence neonatal immune tolerance and microbial interactions ^58^. We observed a significant relationship between serological antibodies and the abundance of specific bacterial taxa in both the mothers and the babies. For instance, in mothers, anti-CBir1 levels were negatively associated with the *Streptococcus* spp. and the *Enterobacteriaceae* family, which are often considered potentially proinflammatory ^59^. Conversely, anti-CBir1 levels were positively associated with *Butyricimonas* spp. and the *Ruminococcaceae* family, which includes essential butyrate producers ^60^. A positive association was also observed with *Bacteroidales,* which are generally beneficial anaerobic bacteria. In babies, cord blood serological biomarkers, including ANCA and anti-OmpC, were linked to lower alpha diversity over time, suggesting that maternal immune responses may negatively influence early-life gut microbiota development. There was also a positive relationship between ANCA and *Romboutsia* spp., a commensal gut inhabitant whose overgrowth could contribute to gut inflammation ^61^. Additionally, ANCA was shown to have significant positive associations with *Citrobacter* spp. and *Pseudomonas* spp., both of which contain opportunistic pathogens ^62,63^, while anti-OmpC also had a positive association with *Pseudomonas* spp. On the other hand, there were positive associations, like one between ASCA IgG level and *Lactobacillus* spp., the abundance of which is considered beneficial ^64^. Given that antibody-microbiota interactions may influence gut barrier function and immune tolerance, future studies should explore how specific antibody-microbiota signatures during early life inform the priming of the immune system in the offspring of IBD patients.

This is the first study that integrates cord blood antibody levels with serial FC and gut microbiota profiling in both mothers and babies, to identify potential inflammatory signals of disease transmission from pregnant women with IBD. However, several limitations must be acknowledged. While the FC and gut microbiota samples were collected throughout pregnancy and during the first three years of life, no peripheral blood was available to assess the correlation with cord blood antibody levels in mothers and babies or to verify the familial concordance of these serological markers, as has been demonstrated in previous work ^65^. Also, despite the prospective study design, the identified associations do not establish causal relationships. Moreover, modest sample size prevented us from conducting additional subgroup analyses, even though the magnitudes of effect that we detected were often meaningful. Our results should be validated in larger, more diverse cohorts to allow a better understanding of the effects of medications and other prenatal risk factors on IBD transmission. Lastly, potential confounders, such as gestational age, maternal disease activity, maternal diet, other health conditions, and environmental exposures, could affect cord blood antibody transfer and should be incorporated in future studies.

## Conclusions

In summary, our findings suggest that maternal IBD-associated antibodies cross the placenta barrier and may contribute to intestinal inflammation and imbalanced microbiota colonization during early life. Whether these serological profiles negatively influence the priming of the baby’s immune system or increase IBD risk in the offspring remains to be determined.

## Acknowledgements

The authors would like to thank all study participants, recruiting physicians, as well as CALPRO AS for providing ELISA kits.

## Disclosure

### Ethics approval and consent to participate

This study was approved by the Institutional Review Board of the Icahn School of Medicine at Mount Sinai (HS#14-00554) as well as the Joint Chinese University of Hong Kong - New Territories East Cluster Clinical Research Ethics Committee (2016.547). This research complies with ethics regulations and infants’ written informed consent were provided by their parents.

### Consent for publication

All patient data included in this manuscript has been de-identified, and therefore, this section is not applicable.

### Data sharing statement

Data from this study can be obtained by making a reasonable request to the corresponding author.

### Authors’ contributions

Conceptualization: IP, JFC, SCN; Methodology: IP, SCN, LZ, DPBM, RD, CY, LT; Investigation: TK, PN; Resources: MP, SCN, DPBM; Data curation: TK, IP, DPBM; Visualization: TK; Writing and editing: TK, IP, PN, MME, AD, IN, AS, LT, JFC, MA, JT, JS; Funding acquisition: JT, SCN, JFC, MA, IP

## Supplementary Information

The online version contains supplementary material.

## Abbreviations

AMOVA: Analysis of Molecular Variance
ANCA: Anti-neutrophil cytoplasmic antibody
ANOSIM: Analysis of Similarities
anti-CBir1: Anti-flagellin CBir1
anti-OmpC: Anti-outer membrane porin C
anti-TNF-α: Anti-tumor necrosis factor-alpha
ASCA: Anti-Saccharomyces cerevisiae antibody
AUCs: Areas Under the Curves
BMI: Body Mass Index
CD: Crohn’s Disease
ELISA: Enzyme-linked Immunosorbent Assay
EU: ELISA Units
FC: Fecal Calprotectin
HBI: Harvey–Bradshaw Index
HK: Hong Kong
IBD: Inflammatory Bowel Disease
IgA: Immunoglobulin A
IgG: Immunoglobulin G
IMM: Immunomodulators
OD: Optical Density
OTUs: Operational Taxonomic Units
PCA: Principal Component Analysis
PCoA: Principal Coordinates Analysis
ROC: Receiver Operating Characteristic
UC: Ulcerative Colitis

