## Supplementary Information for "Maternal IBD-related antibodies are associated with early life gut inflammatory status and microbiota composition: insights from cord blood"

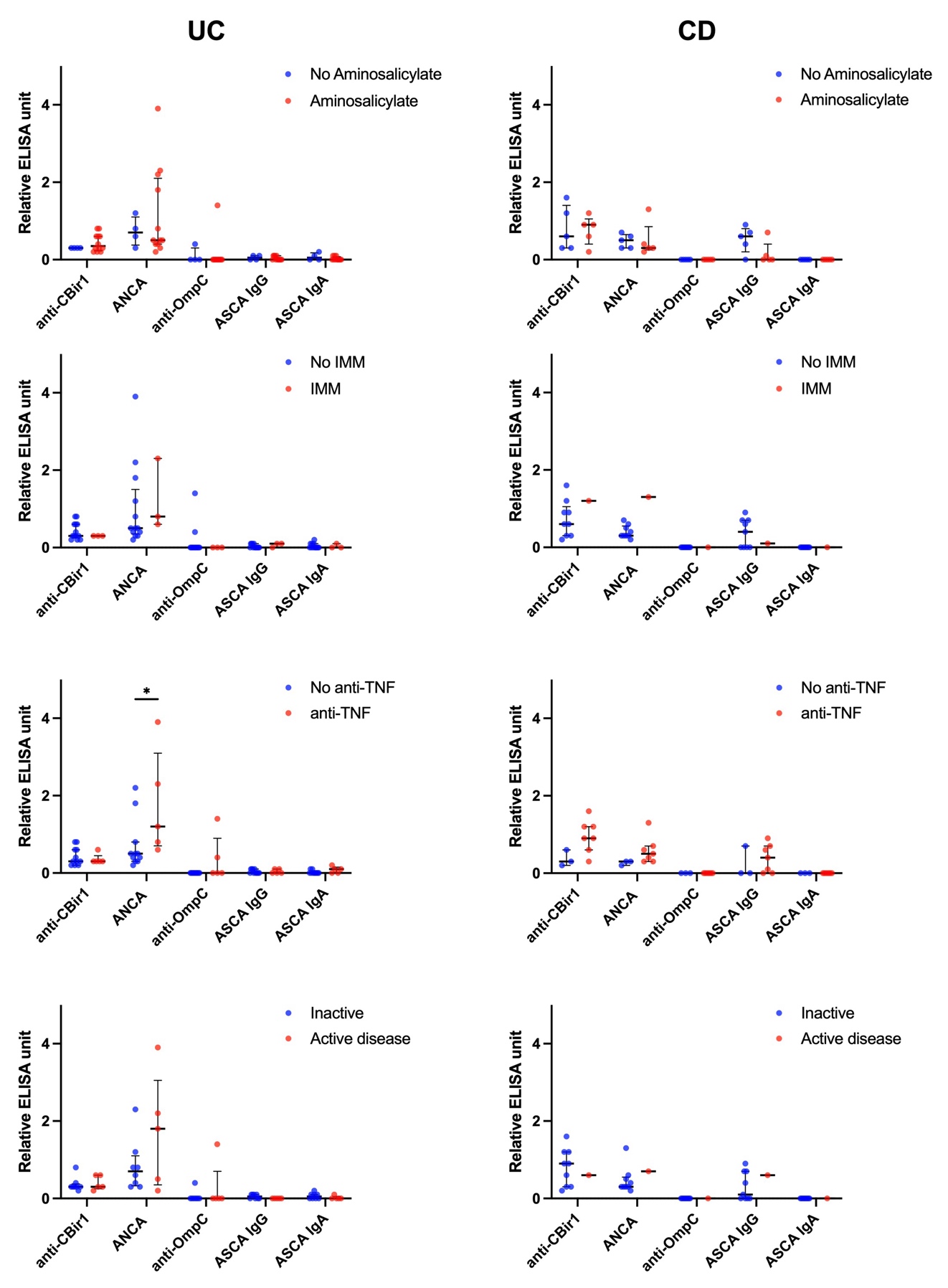

### **Figure S1.** Relative ELISA units of five serological biomarker levels in cord blood with respect to the seropositivity threshold used.

Comparisons were made within ulcerative colitis (UC) or Crohn’s disease (CD) patients by drug treatment (IMM, immunomodulators, anti-TNF-α, anti-tumor necrosis factor α) or by disease activity during pregnancy, only recorded in the mothers recruited in the US. *p < 0.05.

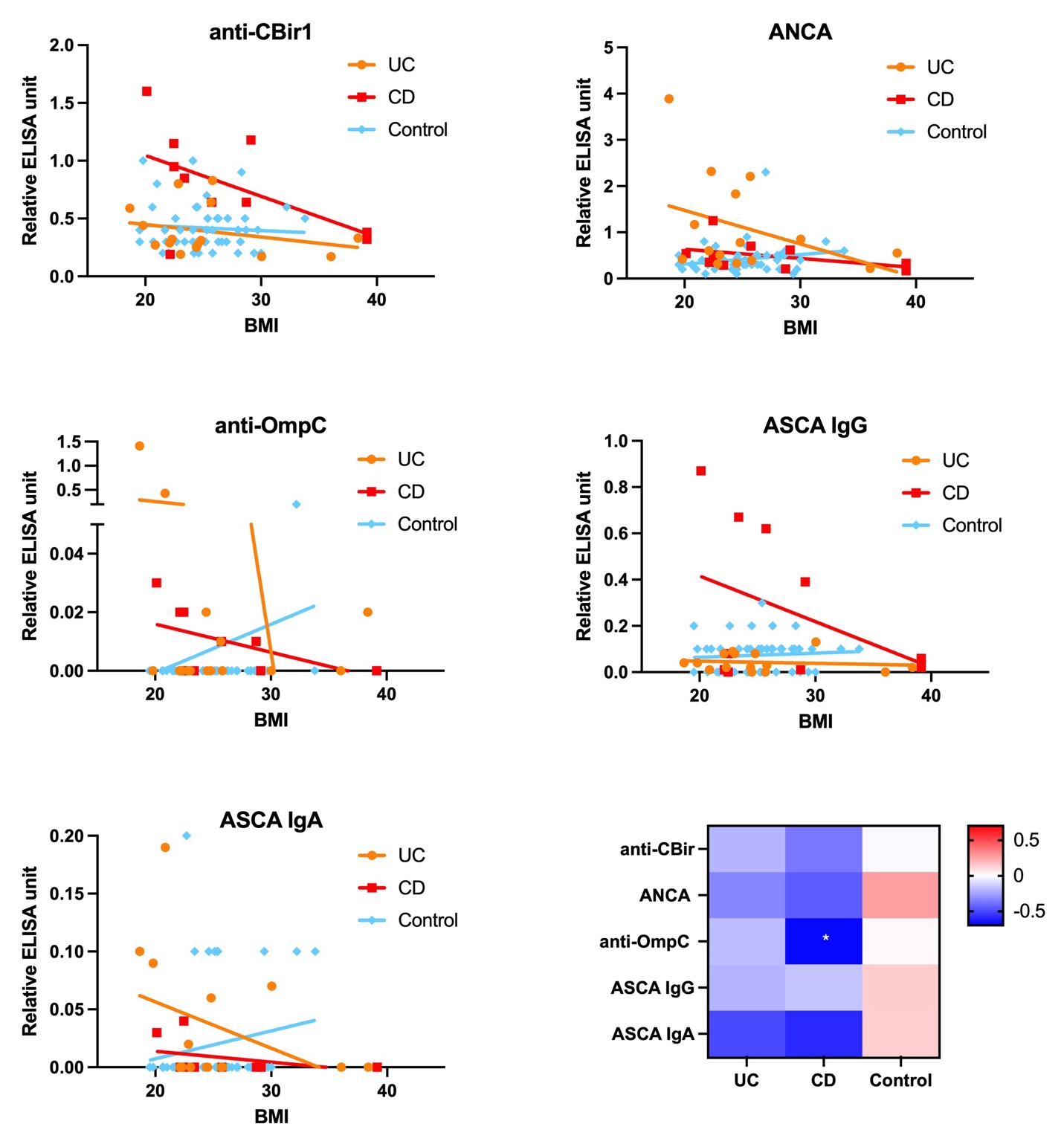

### **Figure S2.** Relative ELISA units of five serological biomarkers in cord blood, plotted against maternal body mass index (BMI) at baseline, with reference to the seropositivity threshold.

Linear regression lines between maternal BMI and each serological biomarker are shown in each panel. The heatmap summarizes Spearman’s correlation coefficients by disease type; *p < 0.05.
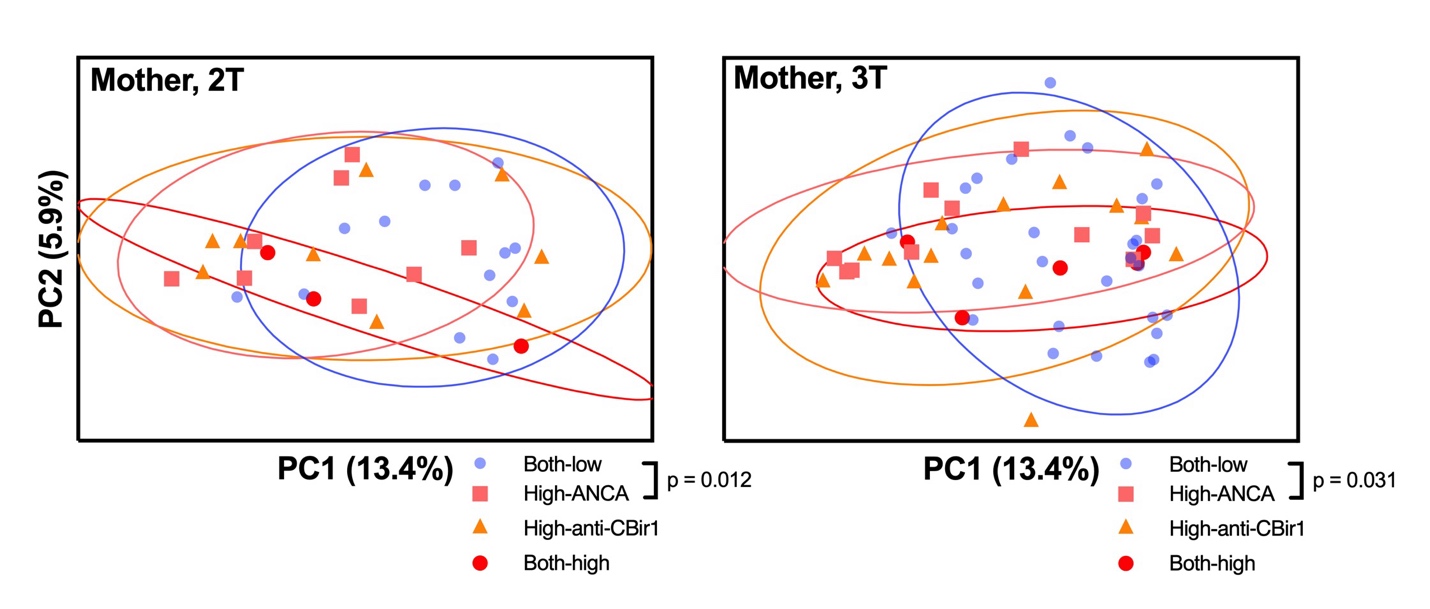

### **Figure S3.** Principal coordinates analysis (PCoA) plot of maternal stool bacterial community composition (beta-diversity), based on Bray-Curtis dissimilarity.

A single PCoA was performed using all maternal samples, and the resulting coordinates are displayed separately for the second trimester (2T) and third trimester (3T) to reduce visual crowding. Ninety percent confidence ellipses are shown. Groups were defined based on the categorical classification (high vs. low) of ANCA and anti-CBir1 in cord blood. Analysis of molecular variance (AMOVA) was conducted at each time point, and statistically significant differences in bacterial composition are indicated with p-values

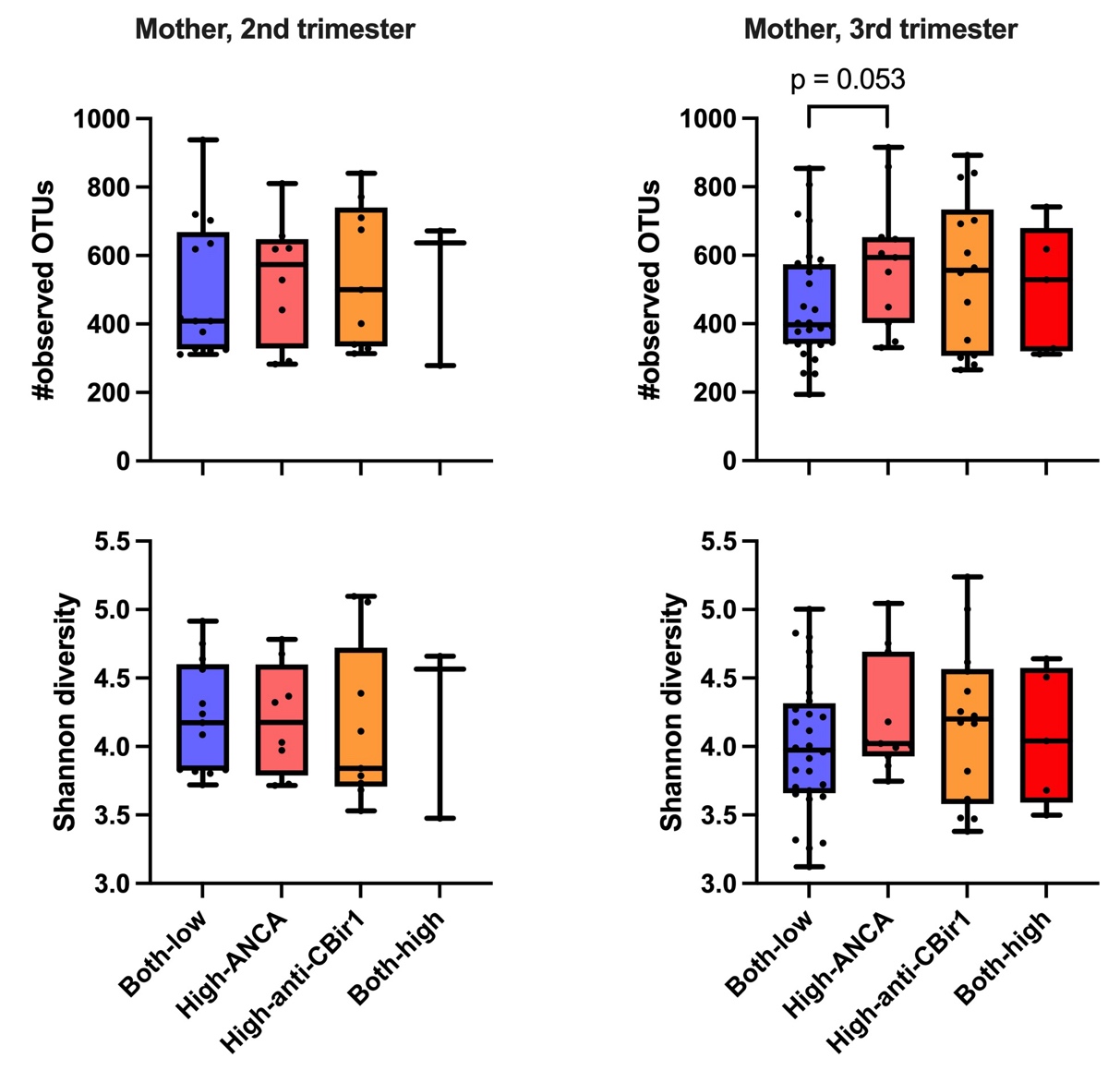

### **Figure S4.** Maternal stool microbiome alpha diversity across four groups defined based on the categorical classification (high vs. low) of ANCA and anti-CBir1 in cord blood.

Second-trimester and third-trimester samples are shown separately. No statistically significant results were observed (p > 0.05); one comparison approached significance (p = 0.053, Wilcoxon rank-sum test).

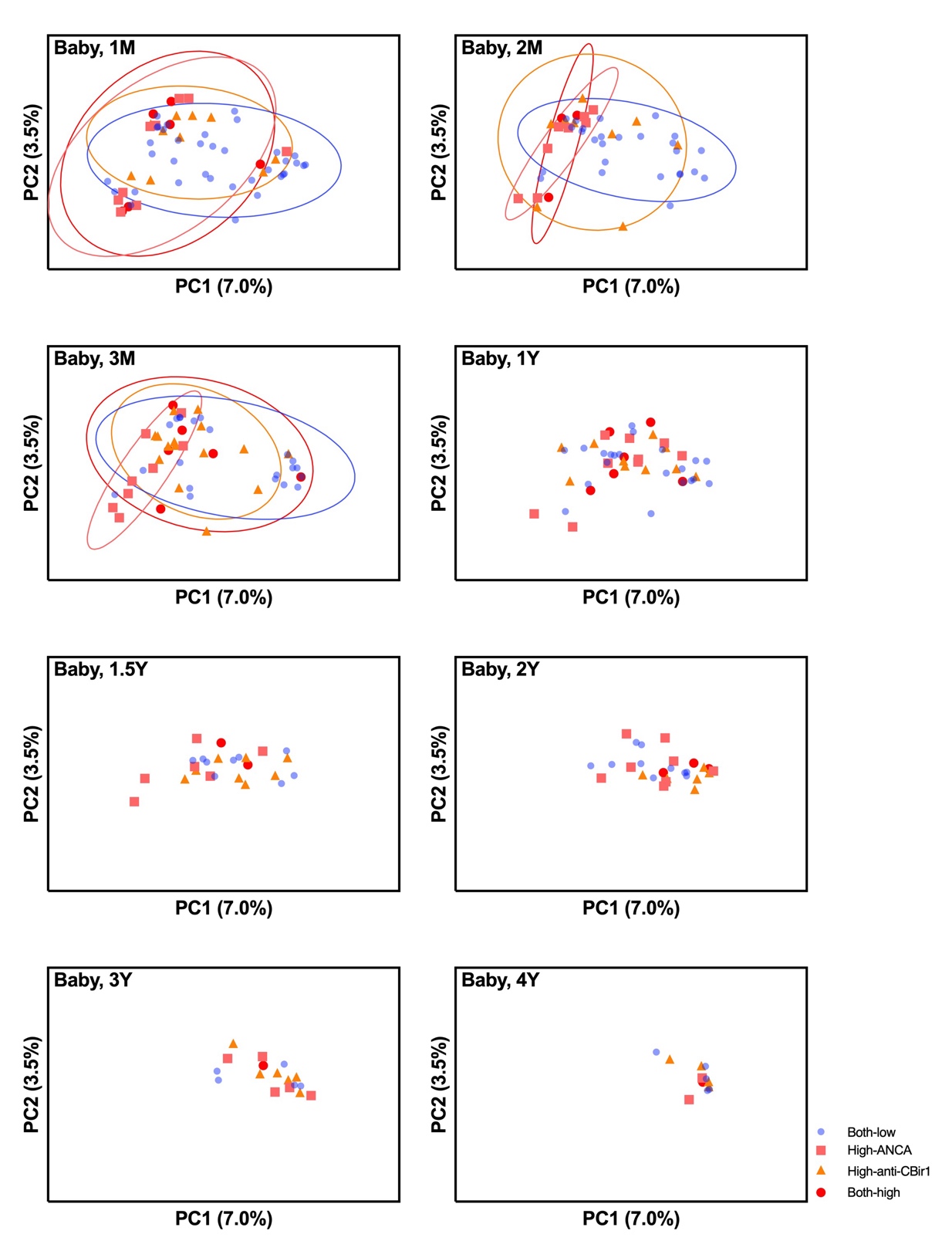

### **Figure S5.** Principal coordinates analysis (PCoA) plot of baby stool bacterial community composition (beta-diversity), based on Bray-Curtis dissimilarity.

A single PCoA was performed using all baby samples, and the resulting coordinates are displayed separately for each time point to reduce visual crowding. Groups were defined based on the categorical classification (high vs. low) of ANCA and anti-CBir1 in cord blood. Analysis of molecular variance (AMOVA) was conducted at each time point among all groups. Ninety percent confidence ellipses are shown for time points where a statistically significant difference was detected. Significant differences were observed between the Both-low and High-ANCA groups at 1 month (p = 0.004), 2 months (p < 0.001), and 3 months (p = 0.003).

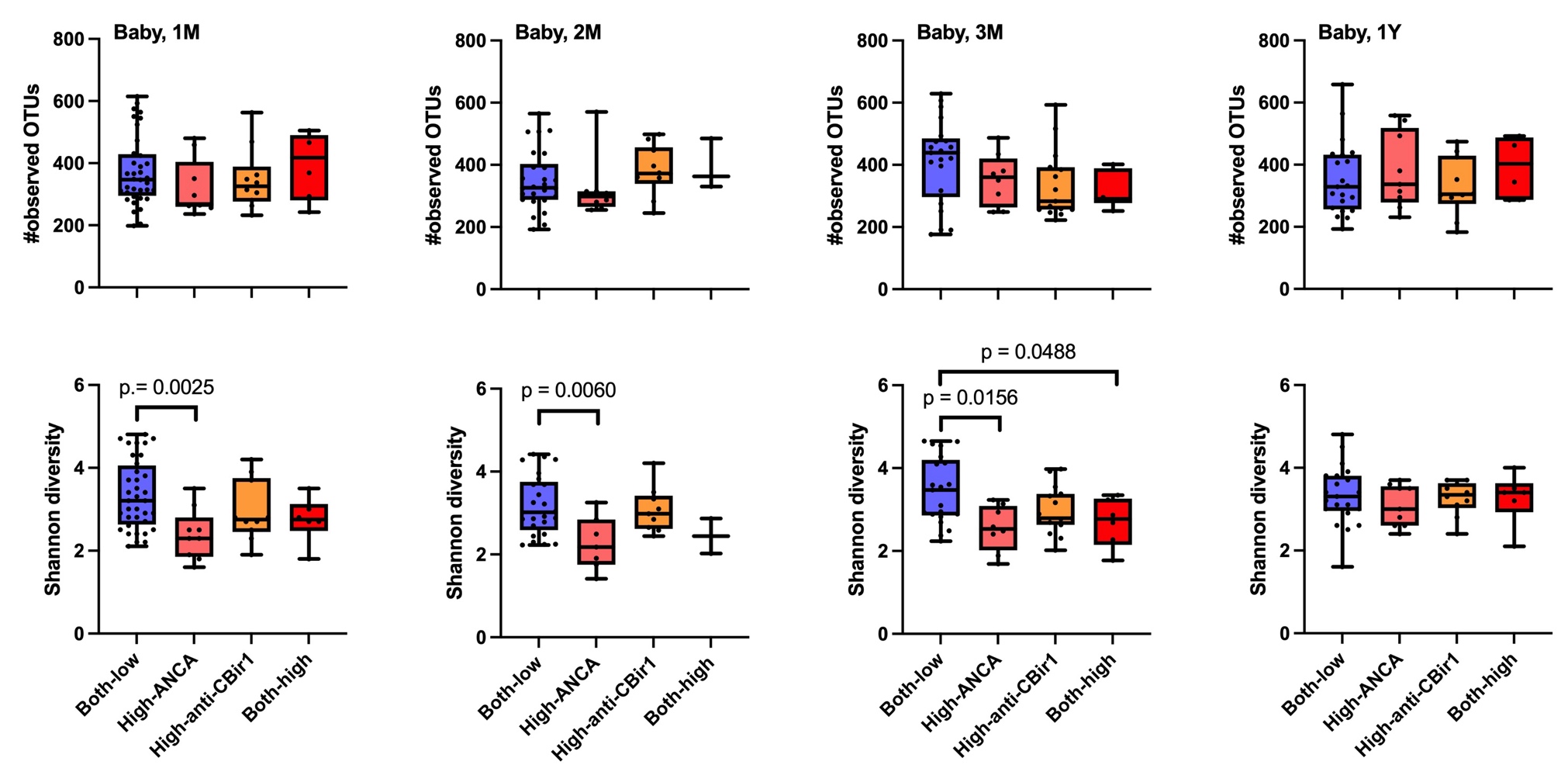

### **Figure S6A.** Baby stool microbiome alpha diversity across four groups defined based on the categorical classification (high vs. low) of ANCA and anti-CBir1 in cord blood.

Four time points (1 month, 2 months, 3 months, and 1 year) are shown separately. Statistically significant results (p < 0.05) by Wilcoxon rank-sum test are shown.

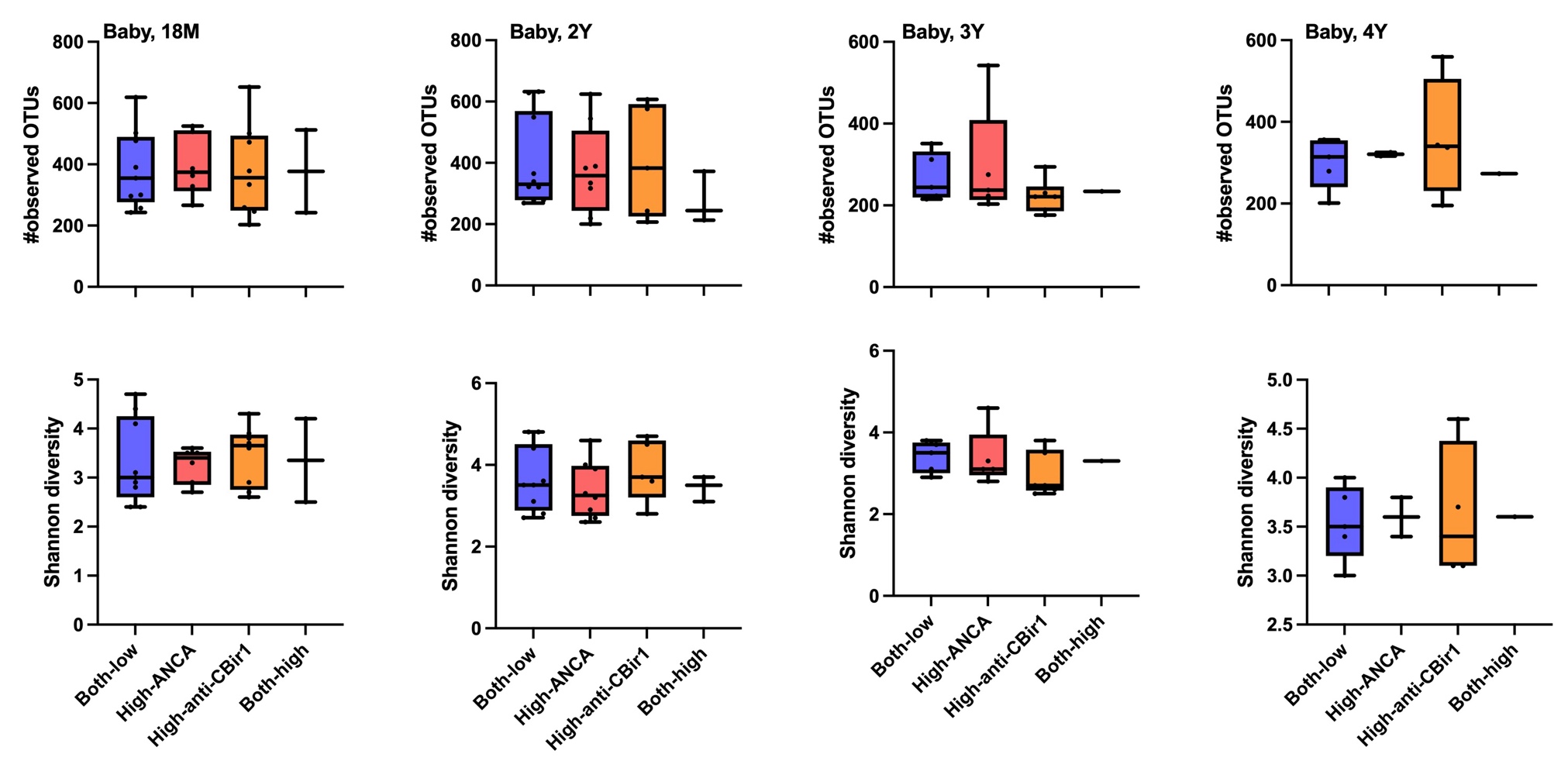

### **Figure S6B.** Baby stool microbiome alpha diversity across four groups defined based on the categorical classification (high vs. low) of ANCA and anti-CBir1 in cord blood.

Four time points (18 month, 24 months, and 36 months) are shown separately. No statistically significant results (all p > 0.05 by Wilcoxon rank-sum test).

### **Table S1.** Combined diagnostic performance of serological markers (anti-CBir1, ANCA, anti-OmpC, ASCA IgG, and ASCA IgA) in cord blood samples

|  | **Sensitivity^1^** | **Specificity^1^** | **Negative predictive value (%)** | **Positive predictive value (%)** | **Accuracy (%)** |
| --- | --- | --- | --- | --- | --- |
| serological marker |  |  |  |  |  |
| UC vs. control | 29.4 | 98.0 | 83.3 | 80.3 | 80.6 |
| CD vs. control | 54.6 | 98.0 | 85.7 | 90.7 | 90.2 |
| UC vs. CD | 88.2 | 72.7 | 83.3 | 80.0 | 82.1 |
| serological marker with BMI |  |  |  |  |  |
| UC vs. control | 33.3 | 98.0 | 83.3 | 83.1 | 83.1 |
| CD vs. control | 70.0 | 98.0 | 87.5 | 94.2 | 93.3 |
| UC vs. CD | 86.7 | 90.0 | 92.9 | 81.8 | 88.0 |

^1^sensitivity and specificity for the first term are shown; sensitivity is the percentage of correctly classified cases when distinguishing ulcerative colitis (UC) from controls, Crohn’s disease (CD) from controls, and UC from CD, respectively.
